# Microbiome structure of ecologically important bioeroding sponges (family Clionaidae): The role of host phylogeny and environmental plasticity

**DOI:** 10.1101/2020.01.28.923250

**Authors:** Oriol Sacristán-Soriano, Xavier Turon, Malcolm Hill

## Abstract

The potential of increased bioerosion by excavating sponges in future environmental scenarios represents a potential threat to coral reef structure and function. If we are to predict changes to coral reef habitats, it is important to understand the biology of these sponges. Little is known about prokaryotic associations in excavating sponges despite the fact that evidence indicates they contribute to the sponge growth through their heterotrophic metabolism and may even act as microborers. Here, we provide the first detailed description of the microbial community of multiple bioeroding sponges from the Clionaidae family (*Cliona varians*, *C. tumula*, *C. delitrix*, *Spheciospongia vesparium*, *Cervicornia cuspidifera*) collected in inshore and offshore coral reefs in the Florida Keys. A total of 6,811 prokaryote OTUs identified using 16S rRNA gene sequencing was detected in the samples studied, including ambient water, belonging to 39 bacterial phyla and 3 archaeal phyla. The microbiomes of species harboring *Symbiodinium* (*Cliona varians*, *C. tumula*, *Cervicornia cuspidifera*) and the azooxanthellate *S. vesparium* were dominated by Alphaproteobacteria that represented from 83 to 96% of total sequences. These clionaid sponges presented species-specific core microbiomes, with 4 OTUs being shared by all sponge samples, albeit with species-specific enrichments. The microbiomes of *C. varians* and *S. vesparium* were stable but showed certain plasticity between offshore and inshore reefs. The distantly related *Cliona delitrix* does not harbor *Symbiodinium,* and had a microbiome dominated by Gammaproteobacteria, which represented 82% of all sequences. Most of the sponge-exclusive OTUs are found in low abundance and belong to the “rare biosphere” category, highlighting the potential importance of these microbes in the ecology of the holobiont. Sponge microbiomes may enhance functional redundancy for the sponge holobiont and allow it to respond to shifting environments over much short time scales than evolutionary change would permit. This work establishes the basis for future research to explore how microbial shifts in bioeroding sponges contribute to bioerosion in the face of a changing environment.

## Introduction

A paradigmatic example of a holobiont is the symbiotic consortium that exists among microbes and their sponge host (Webster and Taylor 2012; Erwin et al. 2015; Thomas et al. 2016; Hill and Sacristán-Soriano 2017; Moitinho-Silva et al. 2017a). Sponges host (even at low relative abundances) up to 60 bacterial and 4 archaeal phyla (Reveillaud et al. 2014; Thomas et al. 2016; Moitinho-Silva et al. 2017a). For most sponges, the within host microbial community is highly diverse and species specific (Thomas et al. 2016). This fact is somewhat surprising given that sponges are filter-feeding bacteriotrophs and thus exposed to a plethora of bacteria from the environment - from transient food items to true sponge associates. It is unclear how sponges discriminate between food items and symbiotic consorts (Hill and Sacristán-Soriano 2017), but it is generally true that sponges sustain a specific microbial composition remarkably different from ambient seawater (e.g., Enticknap et al. 2006; Schmitt et al. 2007; Sharp et al. 2007; Schmitt et al. 2011; Thomas et al. 2016; Turon et al. 2018; Sacristán-Soriano et al. 2019).

The composition of the symbiotic community within sponges is generally host-specific and not a random sample of microbes from the environment (e.g., Hill et al. 2006; Erwin et al. 2012; Schmitt et al. 2012; Pita et al. 2013; Erwin et al. 2015; Steinert et al. 2016; Hill and Sacristán-Soriano 2017; Sacristán-Soriano et al. 2019). Indeed, these associations appear to be consistent over different geographical regions and under different environmental conditions (Hentschel et al. 2002, 2006; Montalvo and Hill 2011; Burgsdorf et al. 2014; Turon et al. 2019). In recent years, high-throughput sequencing methods have generated an extraordinary amount of information on the characterization and functional diversity of associated microbial communities (Hill and Sacristán-Soriano 2017). The perception of the specificity of sponge-associated microbes has changed, and several bacterial taxa thought to be specific to sponges have been shown to occur also in other habitats, such as seawater, sediment and other hosts (Simister et al. 2012). Over 40% of the 173 previously described “sponge-specific” clusters have been detected in seawater (Taylor et al. 2013). So, we may rather use the terms ‘sponge-enriched’ or ‘host-enriched’ to refer to the associated microbial consortia (Moitinho-Silva et al. 2014). Microorganisms that make up the symbiotic community make valuable contributions to many aspects of the sponge physiology and ecology (Taylor et al. 2007). Evidence indicates the symbionts promote the growth and development of the host through the production of regulatory signaling molecules, antibiotics, active secondary metabolites, nutritional components, and other important compounds (Hentschel et al. 2006; Taylor et al. 2007; Webster and Thomas 2016). The holobiont should be a focus of study because organismal phenotype is an integrated product from host and symbiont that shapes all observed benthic marine habitats (Bell 2008). This may be especially true for symbioses in tropical coral reefs that have high rates of productivity despite low availability of environmental inorganic nutrients (Muscatine and Porter 1977; Yellowlees et al. 2008).

Excavating sponges play important ecological roles in nutrient cycling and in sculpting the three-dimensional structure of coral reefs (Rützler 2012; de Goeij et al. 2013; Schönberg et al. 2017a). Bioeroding sponges often account for 40 up to 90% of reef macroborer activity (Schönberg et al. 2017a). Many excavating sponges in the family Clionaidae host photosynthetic dinoflagellates (family Symbiodiniaceae) that help penetrate calcium carbonate reef structures by providing energy to the sponge (Hill 1996; Fang et al. 2014; Achlatis et al. 2018; Achlatis et al. 2019). It has been documented that sponge bioerosion may be enhanced by ocean warming, acidification and eutrophication irrespective of the presence of photosymbionts (Fang et al. 2013; DeCarlo et al. 2015; Silbiger et al. 2016; Schönberg et al. 2017a, b) but with certain physiological constraints (Achlatis et al. 2017). The potential of increased bioerosion by excavating sponges in future scenarios is a threat to coral reefs that deserves greater attention. Most research on bioerosion relates the sponge performance with the activity of their photosynthetic dinoflagellates (e.g., Hill 1996; Weisz et al. 2010; Fang et al. 2014; Achlatis et al. 2018). However, to fully understand bioerosion caused by sponges, we must understand all components of the holobiont, including the prokaryotes, which may influence the growth of sponges through their heterotrophic metabolism. They may also act as microborers themselves, as Schönberg et al. (2019) found evidence of traces of microbial bioerosion in coral cores simultaneously active with the sponge bioerosion.

In the present study, we assessed and compared prokaryote communities from five sponge species belonging to the Clionaidae family from the Florida Keys, FL, USA. Three of the species harbor *Symbiodinium* populations, and two do not. Two of the species are habitat generalists and occur in deep and shallow habitats. We used a culture-independent characterization of microbial communities found in sponges and surrounding seawater using high throughput sequencing of the16S rRNA gene (V4 region). Here, we provide the first detailed description of the microbial community of multiple bioeroding sponges. We sought to answer the following questions: 1) What is the diversity and microbial community composition associated to tropical Clionaidae sponges, compared to the surrounding seawater and with regard to the presence of dinoflagellate symbionts? 2) Is there a core-microbiome associated to them? 3) Are these communities host-specific or do they vary between offshore and inshore reefs?

## Methods and materials

### Sample collection

On May 2017, five sponge species belonging to the Clionaidae family were collected at two habitats in the Florida Keys (USA, FL; Table 1). Among the differential characteristics between the two habitats, we found a wide-range in the thermal regime (from 27 to 34°C during summer months) and variable pH conditions (8.0 to 8.2), with marked tides at the inshore reef (personal observation). Replicate seawater samples (n = 3, l l samples) were collected in sterilized bottles adjacent to the sampled sponges in the field from the offshore and inshore reefs. Sponges were transported to the lab where they were processed within 0.5 to 1 hour of collection. A sample from each sponge was taken with a sterile scalpel and rinsed several times in 0.22 µm-filtered seawater to discard loosely attached microorganisms. Seawater samples were sequentially passed through polycarbonate 5 µm and 0.22 µm filters (MilliporeSigma, Burlington, MA, USA), and the contents on the 0.22 µm filters were used to examine the ambient bacterioplankton communities. All samples were snap-freezed in liquid nitrogen until processed.

**Table 1.**
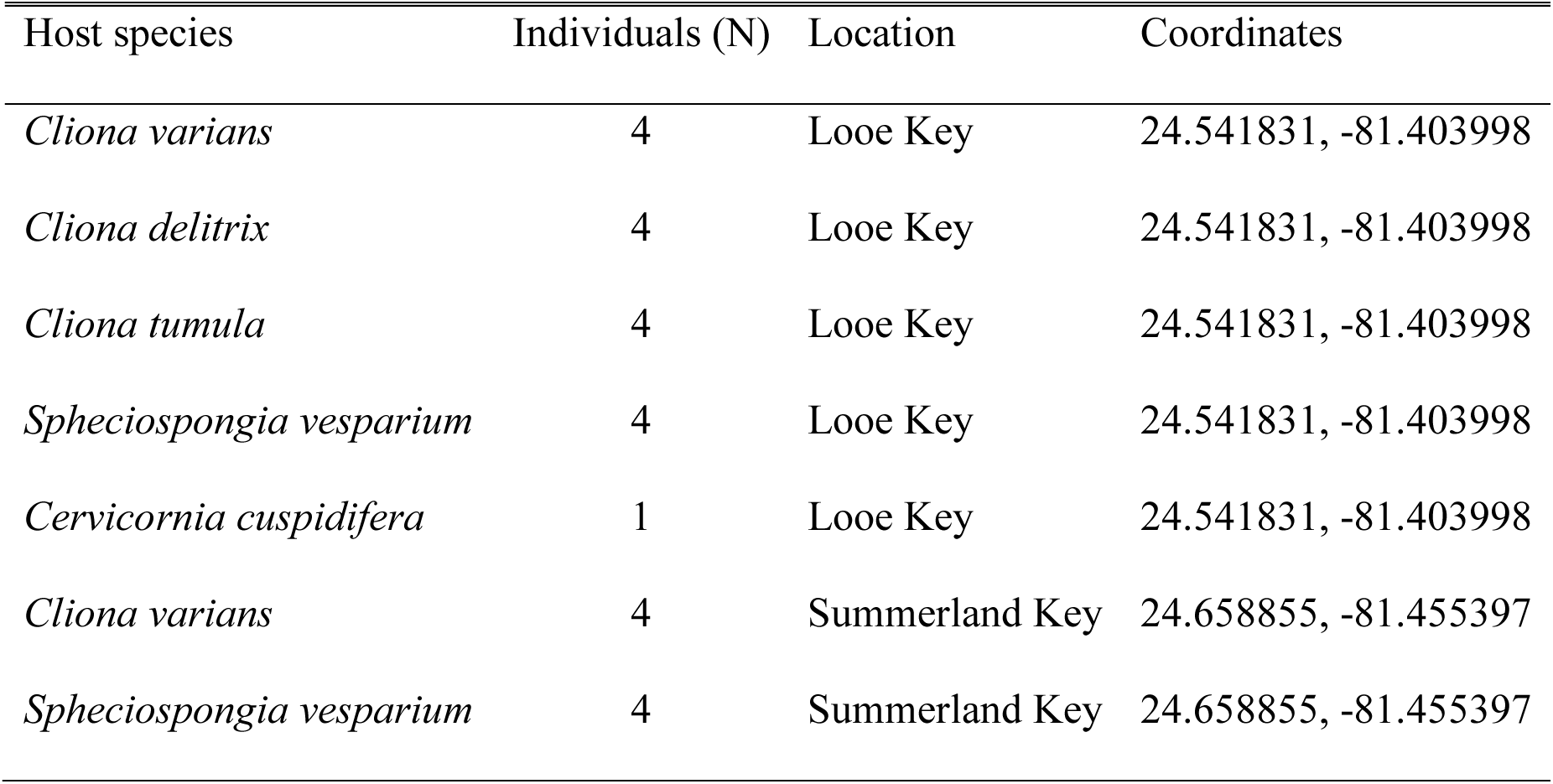
Samples of healthy specimens of *Cliona varians* (Duchassaing & Michelotti, 1864), *Cliona delitrix* (Pang, 1973), *Cliona tumula* (Friday, Poppell & Hill, 2013), *Spheciospongia vesparium* (Lamarck, 1815), and *Cervicornia cuspidifera* (Lamarck, 1815) collected at two offshore (10-12 m deep) and inshore (0.5-1 m deep) reefs in Florida Keys (USA, FL).

| Host species | Individuals (N) | Location | Coordinates |
| --- | --- | --- | --- |
| *Cliona varians* | 4 | Looe Key | 24.541831, -81.403998 |
| *Cliona delitrix* | 4 | Looe Key | 24.541831, -81.403998 |
| *Cliona tumula* | 4 | Looe Key | 24.541831, -81.403998 |
| *Spheciospongia vesparium* | 4 | Looe Key | 24.541831, -81.403998 |
| *Cervicornia cuspidifera* | 1 | Looe Key | 24.541831, -81.403998 |
| *Cliona varians* | 4 | Summerland Key | 24.658855, -81.455397 |
| *Spheciospongia vesparium* | 4 | Summerland Key | 24.658855, -81.455397 |

### Microbiome analysis

DNA was extracted using the DNeasy PowerSoil kit (QIAGEN, Germantown, MD, USA) following standard protocols of the Earth Microbiome Project (http://press.igsb.anl.gov/earthmicrobiome/emp-standard-protocols/dna-extraction-protocol/). DNA extracts were sent to Molecular Research LP (www.mrdnalab.com, Shallowater, TX, USA) for amplification, library construction and multiplexed sequencing of partial (V4) 16S rRNA gene sequences on an Illumina MiSeq platform. The HotStarTaq Plus Master Mix kit (Qiagen) was used for PCR amplifications using DNA extracts as templates with universal bacterial/archaeal forward and reverse primers 515fb (5’-GTGYCAGCMGCCGCGGTAA-3’) and 806rb (5’-GGACTACNVGGGTWTCTAAT-3’), respectively (Caporaso et al. 2011; Apprill et al. 2015). To barcode samples, a multiplex identifier barcode was attached to the forward primer. The thermocycler profile consisted of an initial denaturation step at 94 °C for 3 min; 28 cycles of 94°C for 30s, 53°C for 40s, and 72°C for 1 min with a final elongation step at 72°C for 5 min. Equimolar concentrations of samples were pooled and purified using Agencourt Ampure XP beads (Beckman Coulter) to prepare DNA library by following Illumina TruSeq DNA library preparation protocol. Sequencing was then performed according to manufacturer’s guidelines on an Illumina MiSeq. Illumina sequence data were deposited in NCBI SRA under the project ID PRJNA590868.

Illumina sequence reads were processed in mothur v1.39.5 (Schloss et al. 2009) as previously described (Thomas et al. 2016). Briefly, forward and reverse reads were assembled, demultiplexed, and sequences <200bp and with ambiguous base calls were removed. Sequences were aligned to the SILVA database (release 128, non-redundant, mothur-formatted), trimmed to the V4 region, and screened for chimeras and errors. A naïve Bayesian classifier and Greengenes taxonomy (August 2013 release, mothur-formatted) was used to aid in the removal of non-target sequences (e.g., chloroplasts, mitochondria). We used the SILVA database (release 132, non-redundant, mothur-formatted) for final taxonomic assignment. The resulting high-quality sequences were clustered into operational taxonomic units (OTUs) defined by clustering at 3% divergence and singletons were removed. We used rarefaction curves (mothur v1.39.5) to plot the OTUs observed as a function of sequencing depth. To avoid artifacts of varied sampling depth on subsequent diversity calculations, each sequence dataset was subsampled to the lowest read count (mothur v1.39.5). To place the obtained OTUs into a wider context, these were compared to the database of the sponge EMP project (Moitinho-Silva et al. 2017a) using local BLAST searches (NCBI-BLAST-2.7.1+).

### Community-level analysis

To compare bacterial community profiles, nonmetric multi-dimensional scaling (nMDS) plots of Bray-Curtis similarity matrices were constructed with mothur (v1.39.5) and R (version 3.4.3; ggplot2 package) from square-root transformed OTU relative abundance data. Species *C. cuspidifera* was removed from subsequent analyses as we had just one replicate (see Results). We also constructed bubble charts in R (version 3.4.3; ggplot2 package) from OTU relative abundances to plot community dissimilarities. Significant differences among sponge species and ambient seawater were assessed using a one-way permutational multivariate analysis of variance (PERMANOVA), with the factor source (all sponge species vs. seawater). Significant differences among sponge species were further assessed with one-way PERMANOVA, with the factor source (*C. varians*, *C. delitrix*, *C. tumula*, and *S. vesparium*). Differences between sponge species and habitats were assessed using a two-way PERMANOVA for the species present in the two habitats, with the factors source (*C. varians* and *S. vesparium*), habitat (offshore vs. inshore) and an interaction term. Pairwise comparisons were subsequently conducted for all significant PERMANOVA results involving factors with more than 2 levels. Permutational multivariate analysis of dispersion (PERMDISP) was used to detect differences in homogeneity (dispersion) among groups for all significant PERMANOVA outcomes. All multivariate statistics were performed using in R (version 3.4.3; with adonis2 and betadisper functions from vegan v2.5-6 package).

We calculated three indices of alpha diversity in mothur v1.39.5 (Schloss et al. 2009) to evaluate community richness and evenness: observed OTU richness, the Simpson index of evenness and the inverse of Simpson index of diversity. One-way analyses of variance (ANOVA) was used to detect differences in diversity metrics among the species from the offshore reef (*C. delitrix*, *C. tumula*, *C. varians*, and *S. vesparium*). Two-way ANOVA was used to detect differences in the species present at both habitats, using the factors source (*C. varians* and *S. vesparium*), habitat (offshore vs. inshore) and an interaction term, followed by pairwise comparisons for any significant factor with more than two levels. All data that did not meet the statistical assumptions was transformed accordingly (log-transformation for inverse of Simpson index). The univariate statistics were performed using R (version 3.4.3; Anova function from car package).

### OTU-level analysis

We analyzed the dataset for patterns in relative abundances of particular OTUs within categories (e.g., sponge vs. seawater, offshore vs. inshore). For this purpose, we removed from the dataset rare OTUs (<0.1% relative abundance) and OTUs with a low incidence across samples (detected in ≤2 samples). We used the Mann-Whitney-U test (or Wilcoxon rank sum test) with FDR p-value correction to identify significantly different patterns in OTU relative abundance among hosts and habitats using QIIME1 (Caporaso et al. 2010). To visualize these differences, we constructed OTU networks with the software Cytoscape v.3.7.2 (Shannon et al. 2003).

## Results

### Microbiome composition associated to Clionaidae sponges

After denoising and filtering our sequence libraries, we obtained a total of 2,002,736 reads with a sample depth ranging from 19,564 to 130,500 reads. As we had 4 replicates per species and location in all cases except for *C. cuspidifera*, we discarded those samples (n = 2) with the lowest number of reads (≤28,657), while keeping at least 3 replicates per sponge and site. To avoid artifacts of sequence depth, we rarefied our libraries to the lowest read count (n = 30,726). The OTU accumulation curves showed a lack of plateau in the samples (Suppl. Fig. S1), which implies that we are not capturing all the richness in the samples but we have recovered the abundant OTUs. Thirty-nine bacterial and 3 archaeal phyla were detected in the 6,811 OTUs recovered from seawater and sponge samples, which were predominantly affiliated to the phyla Proteobacteria and Bacteroidetes (Suppl. Fig S2). Of these, 1,949 OTUs were recovered from *C. varians*, 2,026 OTUs from *S. vesparium*, 2,028 OTUs from *C. delitrix*, 1,468 OTUs from *C. tumula* and 345 OTUs from *C. cuspidifera*. In total, 4,352 OTUs were detected exclusively in the sponge samples, while we recovered 2,459 OTUs from seawater, 580 of which were shared with *C. varians* and 576 with *S. vesparium*. The other reef sponges *C. delitrix*, *C. tumula* and *C. cuspidifera* shared 450, 321 and 171 OTUs, respectively, with the ambient seawater sampled from the offshore reef (Suppl. Fig. S3).

The taxonomic composition of microbial communities recovered from surrounding seawater and sponge hosts was markedly different (Fig. 1). The microbial community harbored by all the sponge hosts sampled was enriched for α-Proteobacteria (>80% of the reads of the microbial community, on average) except for *C. delitrix* that was enriched for γ-Proteobacteria (85% of relative abundance). Seawater instead was dominated by more than one bacterial group, α-Proteobacteria (50%) and γ-Proteobacteria (19%). The composition by number of OTUs (instead of abundance) was more balanced, with less dominance of a single or a few groups (Suppl. Fig S4), *C. delitrix* presented a larger fraction of γ-Proteobacteria and the other hosts showed greater OTU richness of α-Proteobacteria. Differences in free-living microbial communities between the offshore reef and the inshore flat reef lay on the relative abundances of Bacteroidetes (4.6% and 31.7%, respectively), Cyanobacteria (9.5% and 0.07%, respectively), Actinobacteria (6.8% and 0.1%, respectively), and Euryarchaeota (2.9% and 0.1%, respectively). Comparatively, these microbial phyla commonly found in seawater samples were depleted in the sponge species analyzed. On the other hand, other less predominant phyla were enriched in the hosts, such as Thaumarchaeota (1.5%) and Planctomycetes (0.3%), compared to planktonic communities (0.02% and 0.07%, respectively). We found a species-specific enrichment in *C. varians* for δ-Proteobacteria (5.5%) while the relative abundance in the other species and in seawater was below 0.6%.

**Figure 1.**
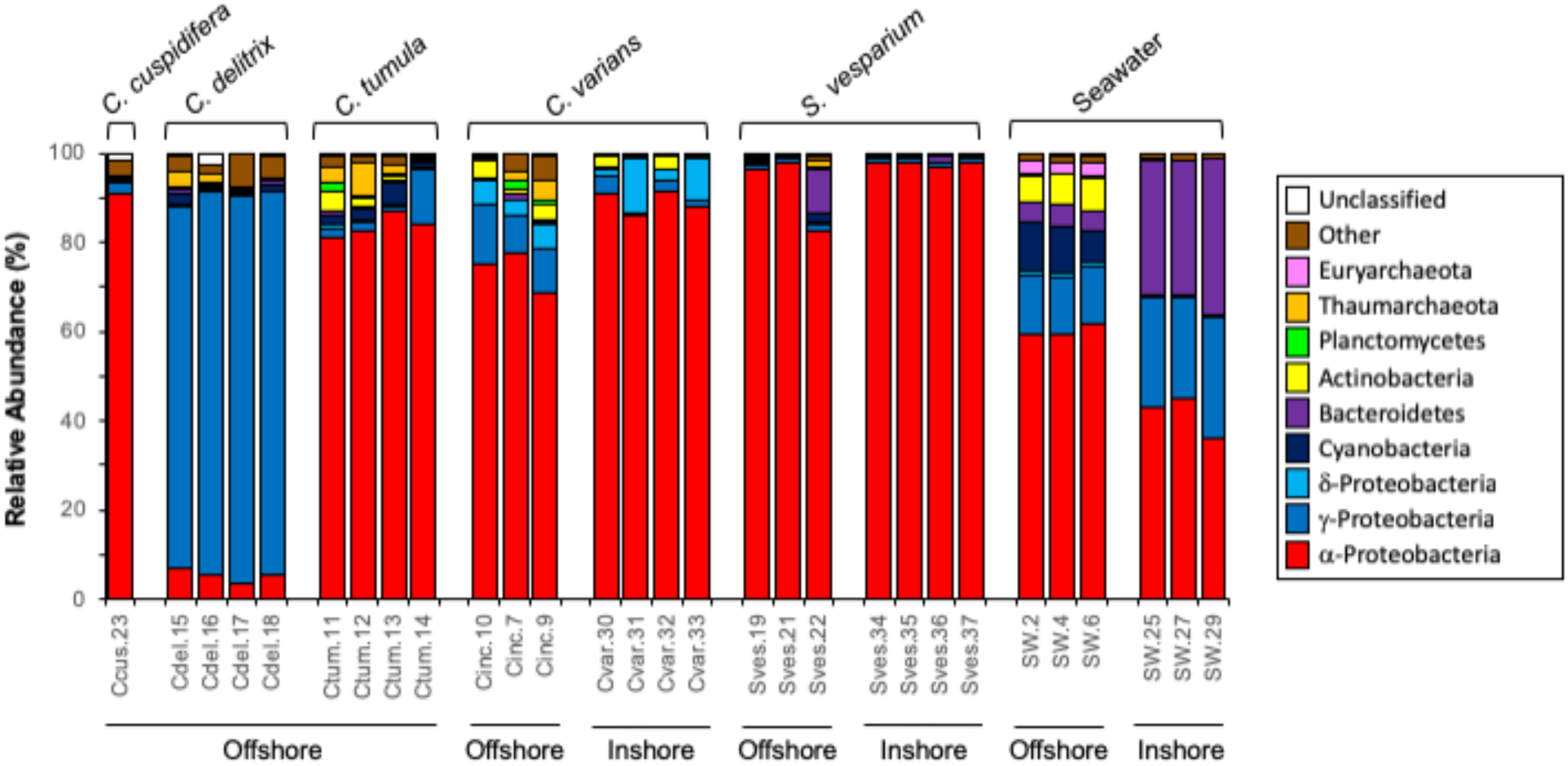
Taxonomic composition of bacterial communities in *Cliona varians*, *Cliona delitrix*, *Cliona tumula*, *Cervicornia cuspidifera*, *Spheciospongia vesparium* and surrounding seawater from Looe Key offshore reef and a Summerland Key inshore reef.

### Differences within and between sponge-associated and seawater microbial communities

Statistically significant differences in microbial community structure (PERMANOVA) were detected among *C. varians*, *C. delitrix*, *C. tumula*, *S. vesparium*, and seawater microbes (F_4,23_ = 6.283; P < 0.001). Symbiont communities from seawater exhibited no overlap with sponge species in the multidimensional space, and all sponge species occupied distinct regions of the nMDS plot (Fig. 2). In addition, a significant interaction between host species (*C. varians* and *S. vesparium*) and habitat occurred (PERMANOVA, F_1,10_ = 2.466; P = 0.031), and thus main factors were analyzed separately. There were significant differences in community structure between offshore and inshore reefs in *C. varians* (t = 4.684, P = 0.026) and *S. vesparium* (t =1.565, P = 0.042). Dispersion analysis revealed equal variability within *C. varians* and *S. vesparium* microbial communities regardless of sampling site (P > 0.05 in all comparisons).

**Figure 2.**
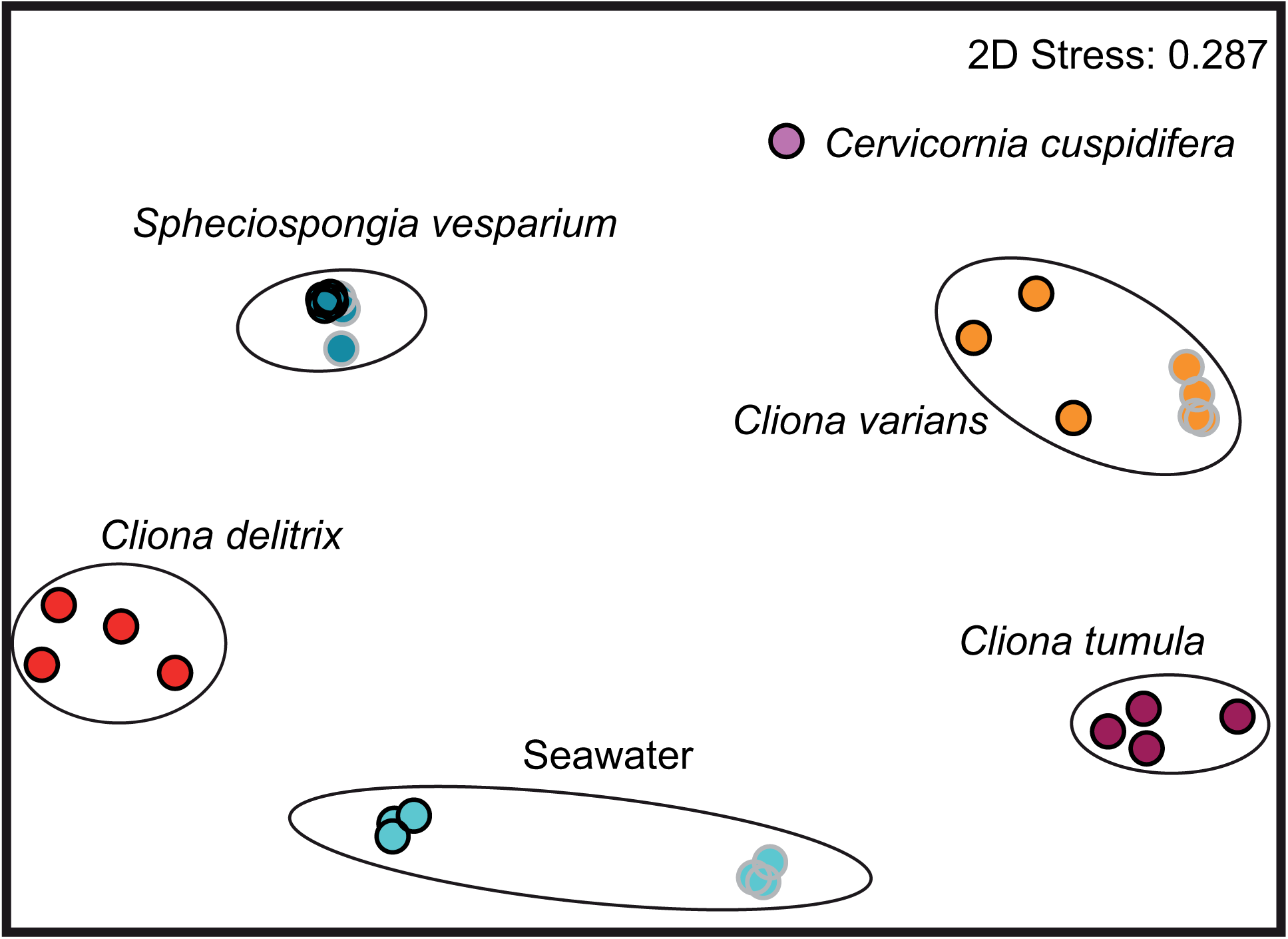
Nonmetric multi-dimensional scaling plot of microbial community structure from replicate individuals of *Cliona varians* (orange), *Spheciospongia vesparium* (dark blue), *Cliona delitrix* (red), *Cliona tumula* (maroon) and surrounding seawater (light blue) from Looe Key (black circles) and Summerland Key (gray circles). Stress value for two-dimensional ordination is shown.

We observed significantly higher mean values of diversity (i.e., inverse Simpson diversity index), and evenness in symbiont communities from seawater compared to host species (P < 0.001 in all pairwise comparisons; Table 2). When we analyzed the sponges from the offshore reef, *C. varians* and *C. delitrix* presented more diverse and even microbial communities than the other species (P < 0.05 in all pairwise comparisons). Comparing *C. varians* and *S. vesparium* from the two habitats studied, a two-way ANOVA detected a significant interaction between hosts and habitats for OTU richness (F_1,10_ = 7.906; P = 0.018) and diversity (F_1,10_ = 9.427; P = 0.012); therefore, main factors were analyzed separately. *C. varians* from the offshore reef harbored richer (P = 0.002) and more diverse (P = 0.002) microbial assemblages compared to the inshore symbiotic community. Considering the community evenness, *C. varians* presented a more even distribution of the microbes hosted compared to *S. vesparium* (F_1,10_ = 25.49; P < 0.001).

**Table 2.**
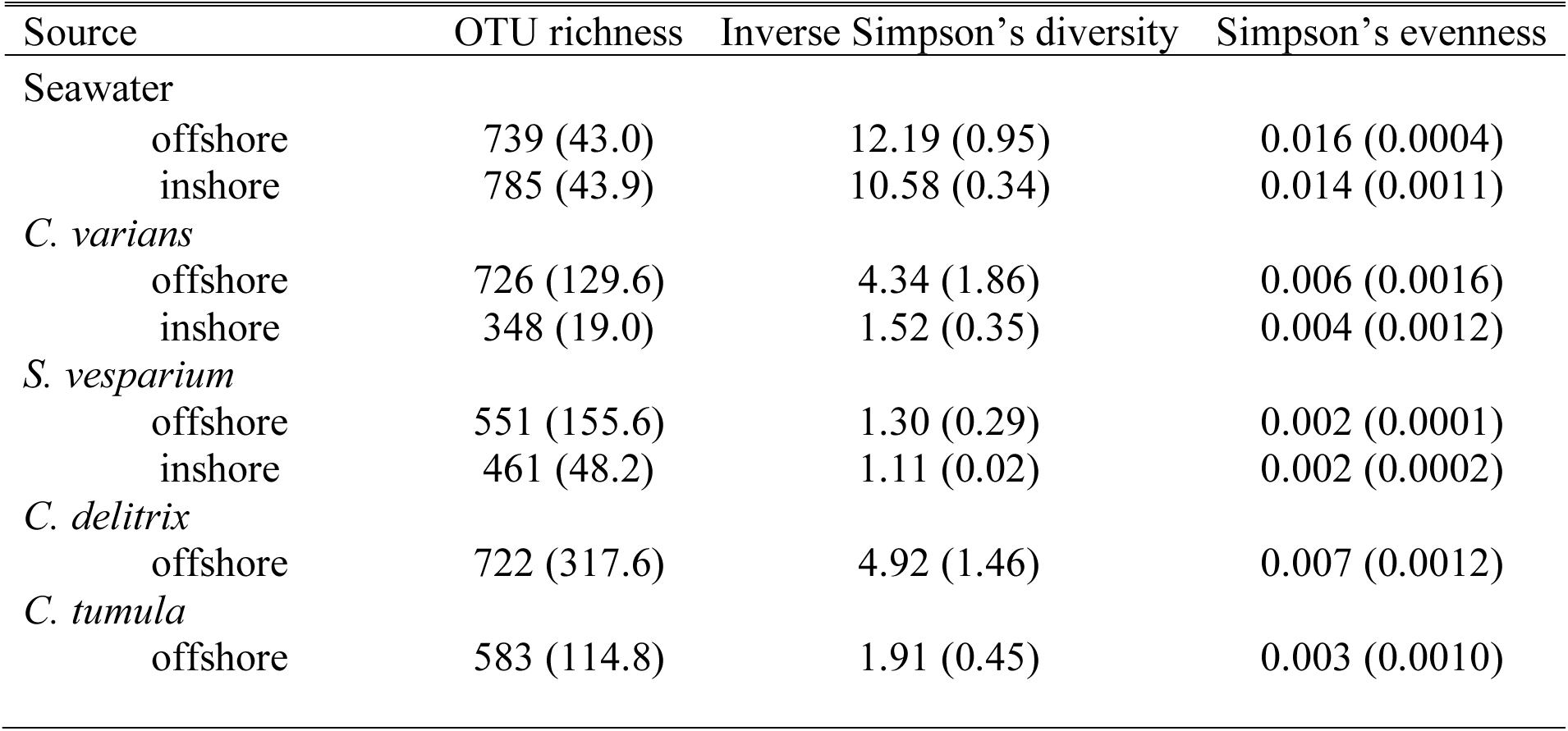
Diversity estimators for microbial communities associated with seawater, *Cliona varians*, *Cliona delitrix, Cliona tumula* and *Spheciospongia vesparium* from Looe Key (offshore) and Summerland Key (inshore). All values represent means (±SE).

The abundance of shared OTUs between sponge-associated and seawater microbial communities was calculated (n = 1,012; 14.9% of the total OTUs recovered) and just 8.3% presented relative abundances over 0.1%. Those few OTUs (n = 84) accounted for 90.6 and 90.3% of the total relative abundance of sponge-associated and seawater microbial assemblages, respectively. All sponge-specific OTUs (n = 4,352; 64% of the total OTUs recovered) fell within the ‘rare biosphere’ (<0.1% relative abundance).

### Core microbiome in sponges from Clionaidae family

In addition to community-level metrics of diversity and structure, we performed a core microbiome analysis to investigate patterns in abundant and prevalent OTUs among sponge hosts. We define here core microbiomes at the species level, as those OTUs shared by all samples of a given species with a mean relative abundance >0.1%. The core microbiome of *C. varians* and *S. vesparium* was formed by 8 and 5 OTUs (Fig. 3A) accounting for over 70% and 90% of total relative read abundance on average, respectively, and 22% and 42% of the number of OTUs with mean relative abundance >0.1%, respectively (Suppl. Table S1). *C. delitrix* and *C. tumula* presented a few more core OTUs (22 and 15, respectively; Fig. 3B) accounting for over 80% of relative abundance in both sponges (65% and 43% of the number of OTUs with mean relative abundance >0.1%, respectively; Suppl. Table S1). Four OTUs (OTU 1, OTU 2, OTU 3 and OTU 36; Fig. 4) were shared by all sponges and were thus present in all defined core microbiomes. The other core OTUs were either shared by two or three species or specific to one host (Fig. 4). Seawater presented a core microbiome of 20 OTUs (including 5 sponge core OTUs) accounting for over 50% of the water microbiome in relative abundance (Suppl. Table S1). We detected significant sponge enrichments in 19 of the 35 sponge core OTUs in at least one of the species analyzed (Suppl. Table S2 for details). *C. varians* was enriched in 5 OTUs (OTUs 2, 15, 30, 44 and 60; cumulative 73% relative abundance) with the predominance of OTU 2 affiliated to Alphaproteobacteria. *C. delitrix* presented an enrichment in 11 OTUs (OTUs 4, 5, 6, 8, 12, 13, 17, 19, 28, 33 and 50; 84% relative abundance) with the dominance of a bacterium assigned to Betaproteobacteriales (OTU 4). *C. tumula* showed a dominant OTU 3 affiliated to Alphaproteobacteria and 3 other enrichments (OTUs 17, 22 and 27; cumulative 81% relative abundance). *S. vesparium* was predominantly enriched in OTU 1, also affiliated to Alphaproteobacteria, and in OTU 60 (accounting for 92% relative abundance; See Fig. 3 & 4). Twenty-seven sponge core OTUs had a mean fold-change in abundance of 100.8 ± 18.8 and were extremely rare in seawater (mean relative abundance <0.01%). Eight additional sponge core OTUs were present with mean relative abundance >2.7% in ambient seawater. From those, 3 OTUs were more abundant in sponges (fold-change 77.5 ± 46.4) and 5 OTUs were enriched in seawater communities (fold-change 26.4 ± 9.7; Suppl. Table S2). If we compared habitats, both *C. varians* and *S. vesparium* presented differences in their core microbiome abundances between offshore and inshore sampling sites (Suppl. Table S2; Fig. 3A).

**Figure 3.**
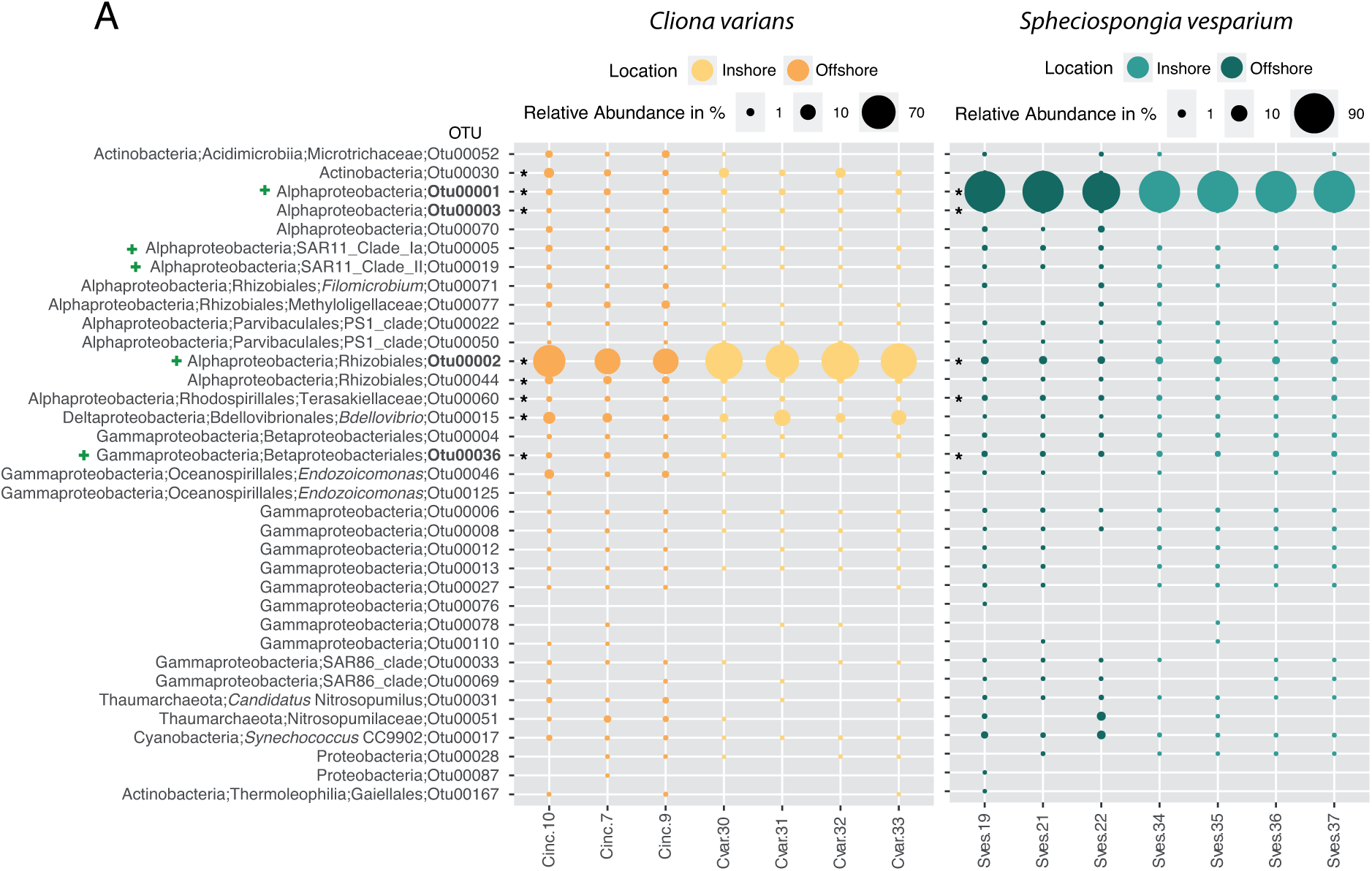
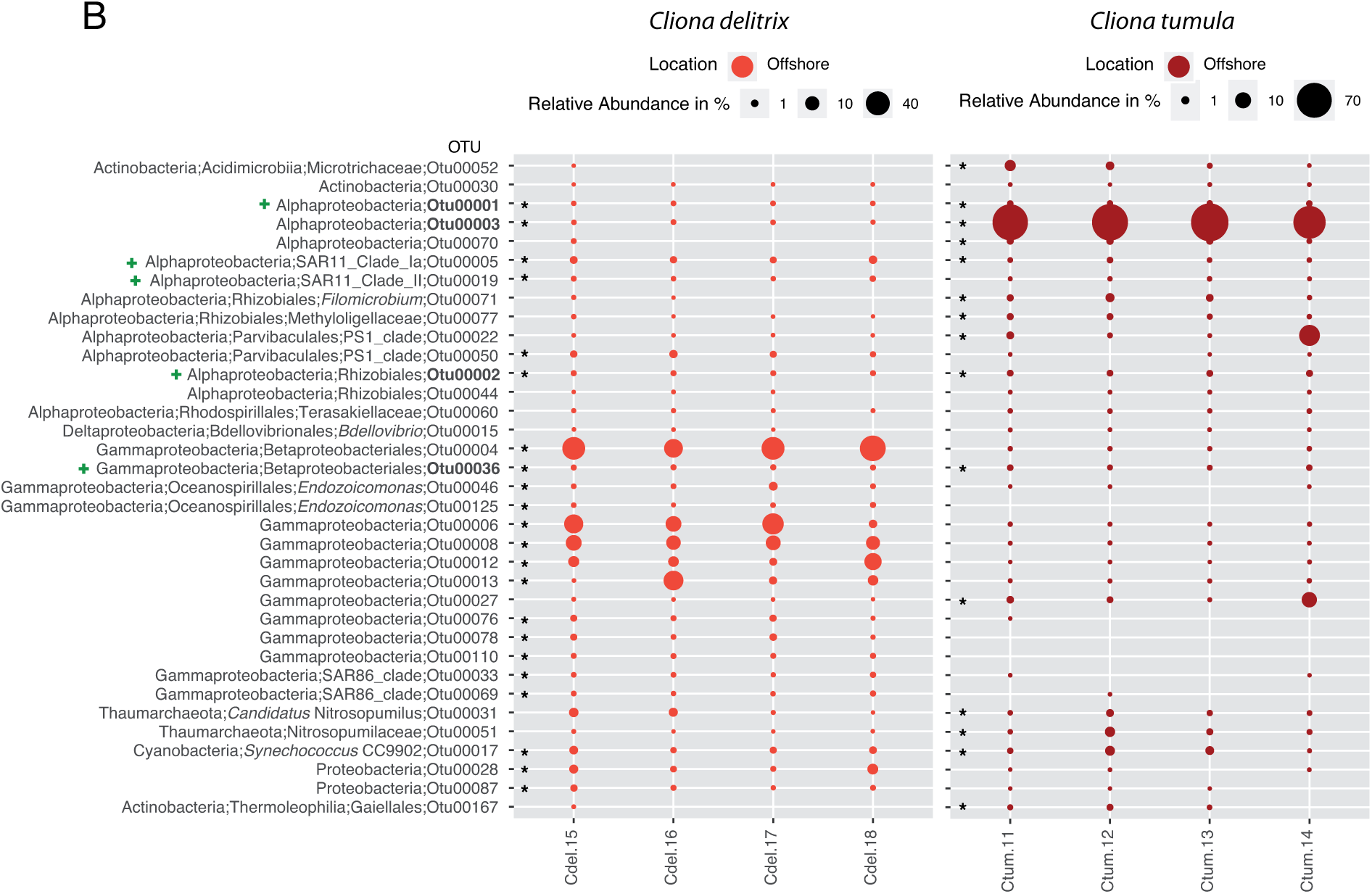
Bubble charts of sponge core OTUs (defined at >0.1% mean relative abundance) of *Cliona varians* - *Spheciospongia vesparium* (A), and *Cliona delitrix* - *Cliona tumula* (B) among habitats. OTU relative abundances are represented by the size of the bubbles (key on the top of each chart; notice the different scales). Asterisks represent the species-specific core microbiome. OTUs shared by the four species are shown in bold. The smallest taxonomical level for each OTU is also shown. Location key: Looe Key reef (Offshore), Summerland Key reef (Inshore). We also show with a green cross those OTUs from core seawater communities.

**Figure 4.**
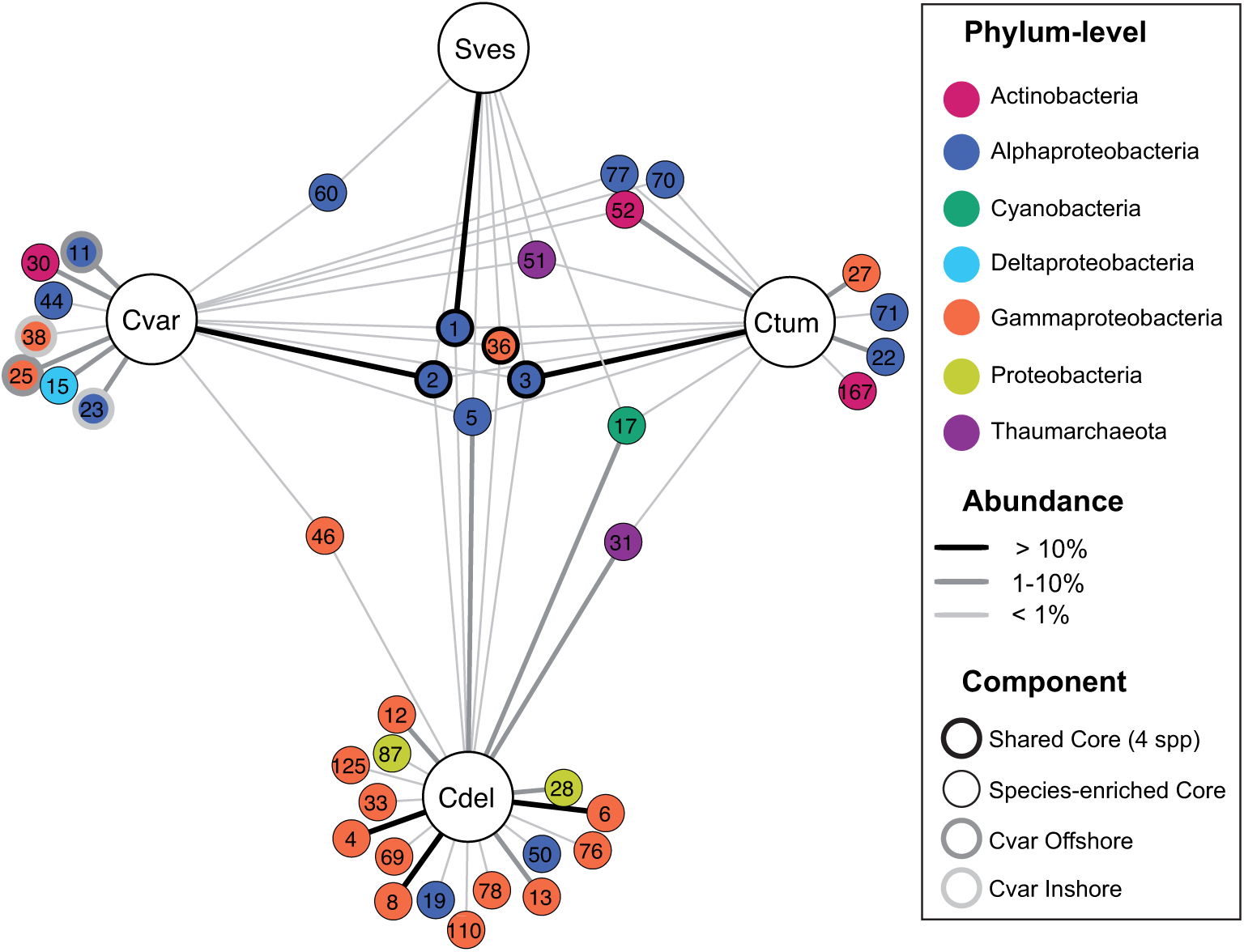
Cytoscape network of the 35 ‘core’ OTUs (present in all replicates and >0.1% abundance) from *C. varians* (Cvar), *C. tumula* (Ctum), *C. delitrix* (Cdel) or *S. vesparium* (Sves). Four other OTUs that differed between inshore and offshore reefs in *C. varians* are also shown. Some OTUs are restricted to specific species whereas others are shared among two, three or the four species analyzed. ‘Core’ OTUs shared by the four species are indicated using bold circle margins. Gray and light gray circle margins indicate OTUs present in Cvar from either offshore or inshore reefs. OTU numbers are shown. Node colors represent the OTU phylum or Proteobacteria class and the edge intensity indicates OTU relative abundance. ‘Rare’ edges (with mean relative abundances <0.1%) were discarded.

### Comparing clionaid associated microbial communities with the sponge EMP database

Local BLAST searches against the sponge EMP database showed that 88% of the OTUs (n = 5960) were found among the sponge microbiome collection with sequence identities over 97%. The core microbiome associated to the sponges from the Clionaidae family is also associated to other sponge hosts and habitats (Suppl. File S1).

## Discussion

This work describes the bacterial and archaeal diversity and the community composition of five sponge species from the Clionaidae family, revealing that the sponges had a microbial signature different from the more diverse and even seawater community. Three of the species studied, *Cliona varians*, *C. tumula* and *Cervicornia cuspidifera*, harbor *Symbiodinium*, whereas *C. delitrix* is free of this dinoflagellate (Hill et al. 2011; Friday et al. 2013; Strehlow et al. 2016). *Spheciospongia vesparium* is not known to harbor *Symbiodinium* and we have not detected this dinoflagellate in our samples under a light microscope (data not shown). The only known species of the genus *Spheciospongia* with *Symbiodinium* cells are *S. inconstans* and *S. vagabunda* (Lévi 1998).

Research on excavating sponges in the last decade is largely focused on estimating bioerosion rates under present and future environmental conditions and determining the role of their photosynthetic symbionts. However, the knowledge of the prokaryotic community associated to bioeroding sponges is limited. Previous research has provided a phylum-level overview of the microbial communities within some *Cliona* species, including *C. celata*, *C. delitrix*, *C. orientalis* and *C. viridis* (Blanquer et al. 2013; Jeong et al. 2015; Pineda et al. 2016; Thomas et al. 2016). In addition, Ramsby et al. (2018) presented detailed species-level community dynamics within *C. orientalis* and how this community responds to seawater warming. Recently, Easson et al. (2020) linked host and microbial genetics on a geographic scale in *C. delitrix*.

### Taxonomic composition and core microbial communities associated to clionaid sponges

Although there is a lot of diversity to be uncovered, we have captured all abundant microbes both in Clionaidae sponges and in seawater (Suppl. Fig. S1). The sponge associated Bacteria/Archaea communities were less diverse than surrounding seawater, reinforcing the view that these sponges were composed of low microbial abundance (LMA) microbiomes, as previously reported for the genus *Cliona* and other clionaids (Poppell et al. 2013; Moitinho-Silva et al. 2017b). Although clionaid sponges displayed less diversity than surrounding seawater, we detected more phyla in sponges (Suppl. Fig S5 & S6). If we discarded those phyla with low sequence abundances (i.e., 0.1% abundance), sponges and seawater harbored 6 bacterial and 1 archaeal phyla. Differences lay in the fact that we detected in sponges groups such as Chlamydiae, Entotheonellaeota and Thaumarchaeota, which were rare in seawater, while we detected Marinomicrobia SAR406, Verrucomicrobia and Euryarchaeota in seawater. However, all those phyla accumulated a microbial abundance ranging from 0.1 to 1.5%. In the case of Archaea, a specific primer pair for this domain might be useful to uncover the archaeal diversity in sponges (Turon and Uriz 2020).

As a general pattern observed in sponge microbiome studies, the phylum Proteobacteria dominates bacterial assemblages, specifically the classes Gamma- and Alphaproteobacteria are the most abundant (Thomas et al. 2016; Moitinho-Silva et al. 2017a; Pita et al. 2018; Cleary et al. 2019). Within the sponge family Clionaidae, Proteobacteria (Gamma- and Alpha- classes) also dominate their microbiomes. However, there is an apparent shift in the class of the dominant Proteobacteria between *Symbiodinium*-bearing and azooxanthellate sponges. *Cliona varians*, *C. tumula* and *Cervicornia cuspidifera* (harboring *Symbiodinium*) were dominated by the class Alphaproteobacteria (from 82.6% to 95.5%), as reported for *C. viridis* and *C. orientalis*, which also present dinoflagellate symbiosis (Blanquer et al. 2013; Pineda et al. 2016; Thomas et al. 2016). *C. delitrix* (a *Symbiodinium*-free species) was instead predominantly occupied by Gammaproteobacteria (85.2% on average), as previously documented for the same species (Thomas et al. 2016; Easson et al. 2020) and for *C. celata* (Jeong et al. 2015), which is categorized as an azooxanthellate species (Miller et al. 2010). However, the microbial composition of *Spheciospongia vesparium* resembled that from *Symbiodinium*-bearing species with dominance of Alphaproteobacteria.

While the presence of *Symbiodinium* may influence the taxonomic composition of the microbiome, it is also important to provide context about the taxonomic challenges presented by the host sponges. Rützler and Hooper (2000) suggested a reorganization of the family Clionaidae to include some sponges that had historically been placed in the family Spirastrellidae. Previously, *C. varians* was in the genus *Anthosigmella*, *C. cuspidifera* was in the genus *Spheciospongia*, and both genera were in the family Spirastrellidae. Rützler and Hooper (2000) moved these species to the Clionaidae based on their capacity to bioerode. Hill et al. (2011) suggested that the taxonomic revision may not have been required given that Clade G *Symbiodinium* appeared to distinguish between ‘spirastrellid-like’ sponges (i.e., *C. varians* and *C. cuspidifera* harbored *Symbiodinium* that had an 86 bp b-loop variant) and true clionaid-like sponges (e.g., *C. orientalis* harbored *Symbiodinium* that had an 85 bp b-loop variant). Thus, an alternative explanation for the patterns we observed in microbiome community composition is that the sponge hosts belong to two distinct poriferan families or phylogenetic clades, and the microbiome differences are driven by host taxonomy and not the presence of *Symbiodinium*. Some studies that have explored the phylogenetic relationships of several species of the family Clionaidae might support this hypothesis. It seems that *C. delitrix* would have evolved earlier and would be distantly related to a well-supported clade formed by *C. varians*, *C. cuspidifera* and three species of the genus *Spheciospongia* (Kober and Nichols 2007; Escobar et al. 2012). If this is true, coevolutionary processes between hosts and their microbial partners appears to play a larger role in shaping microbe community composition than the presence of *Symbiodinium*. Further research is needed to assess the importance of coevolutionary history or the interactions among multiple microbial partners within the sponge in driven microbiome community composition.

The sponges *C. varians* and *S. vesparium* presented 8-5 core components in which nearly 70% and 90% of all 16S rRNA reads belonged to a single OTU, with highest similarity to Rhizobiales (Alphaproteobacteria) and unclassified Alphaproteobacteria, respectively. *C. tumula* and *C. delitrix* harbored instead a more diverse core microbiome (15-22 OTUs), but both hosts also followed the microbial signature of LMA sponges with the dominance of a single OTU in *C. tumula* (72%), ascribed to unclassified Alphaproteobacteria, and a couple of OTUs (36% and 17%) in *C. delitrix*, with highest similarity to Betaproteobacteriales (Gammaproteobacteria) and unclassified Gammaproteobacteria, as recently found for the latter species (Easson et al. 2020). The low resolution of the taxonomic assignment precludes functional analyses and hinders shedding light on the role of symbiotic partners. The dominant OTUs from *C. varians*, *C. tumula* and *S. vesparium* were shared by the other clionaid species, but with relative abundances much lower, ranging from 0.1 to 0.3%. Likewise, the two dominant OTUs from *C. delitrix* were depleted in the other hosts and assigned to the ‘rare biosphere’ (<0.1% reads). Nearly 80% of sponge core components were not found or extremely rare (<0.01% on average) in the surrounding seawater representing a 100-fold increase. From the remaining 20%, three core OTUs were 77 times more abundant on average in sponges and five of them showed a 26-fold change in the environment. These results point to a strong selective ability of the sponges, as found in previous studies (Turon et al. 2018).

The low number of core microbial components in *C. varians* and *S. vesparium* are due to microbial differences between offshore and inshore environments. As more locations are sampled of a particular species, the more reduced core microbiome can be detected. These differences were more evident in the former species, where the core OTU 2 was predominant in the inshore specimens (82% vs. 46%). Besides this compositional change between habitats, four other bacterial components were highly common in one of the sites while extremely rare in the other (Fig. 4). Two OTUs were assigned to the Alphaproteobacteria class and the other two were affiliated with the genera *Endozoicomonas* (OTU 25) and *Pseudohongiella* (OTU 38), both from the class Gammaproteobacteria. The genus *Endozoicomonas* is commonly found in close association with sponges (Nishijima et al. 2013) and other invertebrates such as corals (Bourne et al. 2016). Multiple functions related to nutrient acquisition and/or cycling, structuring the sponge microbiome via signaling molecules or roles in host health have been proposed for this genus (Nishijima et al. 2013; Rua et al. 2014; Gardères et al. 2015; Morrow et al. 2015; Neave et al. 2016). The genus *Pseudohongiella* has been frequently reported in marine bacterioplankton (Xu et al. 2019) but has been also found in sponge microbiomes (Chaib De Mares et al. 2018). Its function is unclear but a recent genomic analysis of this genus in pelagic environments reveals adaptation mechanisms to enhance abilities in the transfer and metabolism of organic and inorganic materials and to react quickly to external changes (Xu et al. 2019). In any case, we found an effect of habitat in the two species analyzed, both in the multivariate composition and in the univariate descriptors. However, significant interaction terms indicated that the response is species-specific. These results are in agreement with a recent study that found an spatial component in the variability of microbiomes within the same species (Easson et al. 2020). Further research is required to investigate if those changes represent an adaptation to different environmental regimes.

The abundance and stability of dominant OTUs among clionaid species suggest a close partnership with the host. Lurgi et al. (2019) revealed that sponges of the order Clionaida shared a microbial organization. This result would support the similarities we found in microbial diversity among the core microbiomes of the sponges from the family Clionaidae, with compositional differences driven by host identity (Thomas et al. 2016). However, clionaid sponges exhibited flexibility of microbial partnerships between and within species and across habitats. This microbial plasticity may serve as a mechanism to preserve selected functions among individuals and species, so these taxonomical shifts may enhance functional redundancy. In open microbial systems, like sponges, taxonomic composition seems to be decoupled from functional structure (Louca et al. 2018) contributing to the sponge microbiome resilience. The degree of functional redundancy depends on the environment and the function considered. Important functions may be better buffered against environmental changes by redundant biodiversity in order to guarantee a proper functioning of the system (Jurburg and Salles 2015; Louca et al. 2018). In our case study, the taxonomic variability found in clionaid sponges at both intraspecific and interspecific levels may produce similar metabolic profiles that contribute to the health and survival of the host. We found that the core microbiomes harbor a high fraction of unclassified bacteria at class or order levels. Given the importance of clionaid sponges to reef bioerosion, further research is needed to identify and classify these microbial strains to fully understand their metabolic potential and determine the role of associated prokaryotic organisms on the sponge eroding capabilities.

Less attention has been paid to the ‘rare biosphere’ (<0.1% relative abundance) of the sponge microbiome, a concept originally described for the deep sea (Sogin et al. 2006). This fraction is highly diverse and controlled by host and environmental factors (Reveillaud et al. 2014; Lurgi et al. 2019). Its metabolic potential might be relevant for sponge functioning since it is not uncommon to observe a replacement of dominant taxa by metabolically similar rare species within a few weeks in microbial systems, such as a wastewater treatment plant or experimental bioreactors (Ofiţeru et al. 2010; Fernandez-Gonzalez et al. 2016). These changes can confer a rapid phenotypic response to changing environments, much faster than is possible via evolutionary selection. Therefore, it is likely advantageous and informative to focus on functional traits rather than taxonomy if we are interested in estimating how communities respond to their habitats under the threat of global warming (McGill et al. 2006). Future studies should explore which functions are altered in the sponge holobiont across environmental gradients so that we may predict the effects of a changing environment at the ecosystem level.

In conclusion, we used high throughput sequencing to provide a detailed characterization of the microbiome of sponges from the Clionaidae family. The *Symbiodinium*-bearing species from this study and the closely related *S. vesparium* were dominated by Alphaproteobacteria, while the azooxanthellate and distantly related *C. delitrix* was dominated by Gammaproteobacteria. These clionaids show a species-specific core microbiome with dominant OTUs partly shared among species but with species-specific enrichments. *C. varians* and *S. vesparium* showed variations in their microbiomes between offshore and inshore reefs probably due to an adaptation to different environmental conditions, although this hypothesis needs to be tested. The other question that arises from the present study is about functional redundancy. Is the plasticity or flexibility of sponge microbiomes related to redundant functions? Given the importance of clionaid sponges to reef bioerosion, understanding the functional basis of prokaryotic symbiosis in holobiont performance is essential. Future research should address how microbial shifts in bioeroding sponges affects sponge resilience and performance under climate change scenarios.

## Acknowledgements

We would like to thank Iosune Uriz for her comments and assistance in light microscopy. This work was funded by grants from the National Science Foundation (OCE-1617255 and IOS-1555440 to MH) and in part by the Spanish Government project PopCOmics CTM2017-88080 (MCIU,AEI/FEDER, UE) to XT and the European Union’s Horizon 2020 research and innovation program under the Marie Sklodowska-Curie grant agreement 705464 (“SCOOBA”) to OSS.

## Supplemental Figures, Tables and Files

Figure S1. Rarefaction curves present the relationship between the sampling effort and the microbiome OTU richness in *Cliona varians* forma *incrustans* (Cinc), *Cliona varians* forma *varians* (Cvar), *Cliona delitrix* (Cdel), *Cliona tumula* (Ctum), *Spheciospongia vesparium* (Sves), *Cliona cuspidifera* (Ccus) and ambient seawater (SW).

Figure S2. Total abundance of the microbiome OTUs recovered from sponges and seawater samples as a function of its prevalence and classified by phyla. Log scale in the x-axis. Discontinuous line indicates 10% prevalence.

Figure S3. Venn diagrams showing the unique and shared microbiome OTUs among *Spheciospongia vesparium* and seawater (A) *Cliona varians* and seawater samples (B) and *Cliona delitrix*, *Cliona tumula*, *Cervicornia cuspidifera* and seawater from the offshore reef (C) defined at distance of 0.03 (i.e., 97% similarity). Inshore *S. vesparium* (Sves_F), offshore *S. vesparium* (Sves_R), inshore *C. varians* (Cvar), offshore *C. varians* (Cinc), *C. delitrix* (Cdel), *C. tumula* (Ctum), *C. cuspidifera* (Ccus) and ambient seawater from offshore (SW_R) and inshore (SW_F) reefs.

Figure S4. Taxonomic composition of bacterial communities in *Cliona varians*, *Cliona delitrix*, *Cliona tumula*, *Cervicornia cuspidifera*, *Spheciospongia vesparium* and surrounding seawater from Looe Key offshore reef and a Summerland Key inshore reef. Fraction of OTUs per sample classified by taxonomic group.

Figure S5. Total abundance of the microbiome OTUs recovered from sponge samples as a function of its prevalence and classified by phyla. Log scale in the x-axis. Discontinuous line indicates 10% prevalence.

Figure S6. Total abundance of the microbiome OTUs recovered from seawater samples as a function of its prevalence and classified by phyla. Log scale in the x-axis. Discontinuous line indicates 10% prevalence.

Table S1. Core microbiome defined at 0.1% mean relative abundance. Core abundances across replicates are also shown. OTUs in bold represent those core OTUs shared among species. Sources: *Cliona varians*, *Cliona delitrix*, *Cliona tumula*, *Spheciospongia vesparium* and seawater.

Table S2. Significantly different abundant OTUs in multiple comparisons among sources according to the false discovery rate (FDR) probabilities. Mean sequence count for the corresponding source is provided with colored values representing higher counts than the other sources compared. The taxonomy affiliation of each OTU is also shown. Sources: *Cliona varians*, *Cliona delitrix*, *Cliona tumula*, *Spheciospongia vesparium* and seawater. Representing sponge Core Microbiome OTUs in bold (at 0.1% Relative Abundance). *represents core OTUs of the species analyzed.

File S1. Local blast results of the microbiome OTUs from *Cliona varians*, *Cliona delitrix*, *Cliona tumula*, *Spheciospongia vesparium*, *Cervicornia cuspidifera* and ambient seawater against the Sponge Earth Microbiome Project database. First hit, alignment matches and sequence identities are shown. Percentage of microbiome OTUs above identity thresholds is also shown. Sponge core OTUs are represented in bold.

