## Supplementary figures and images for "Microbiome structure of ecologically important bioeroding sponges (family Clionaidae): The role of host phylogeny and environmental plasticity"

### Supplemental Figure 1

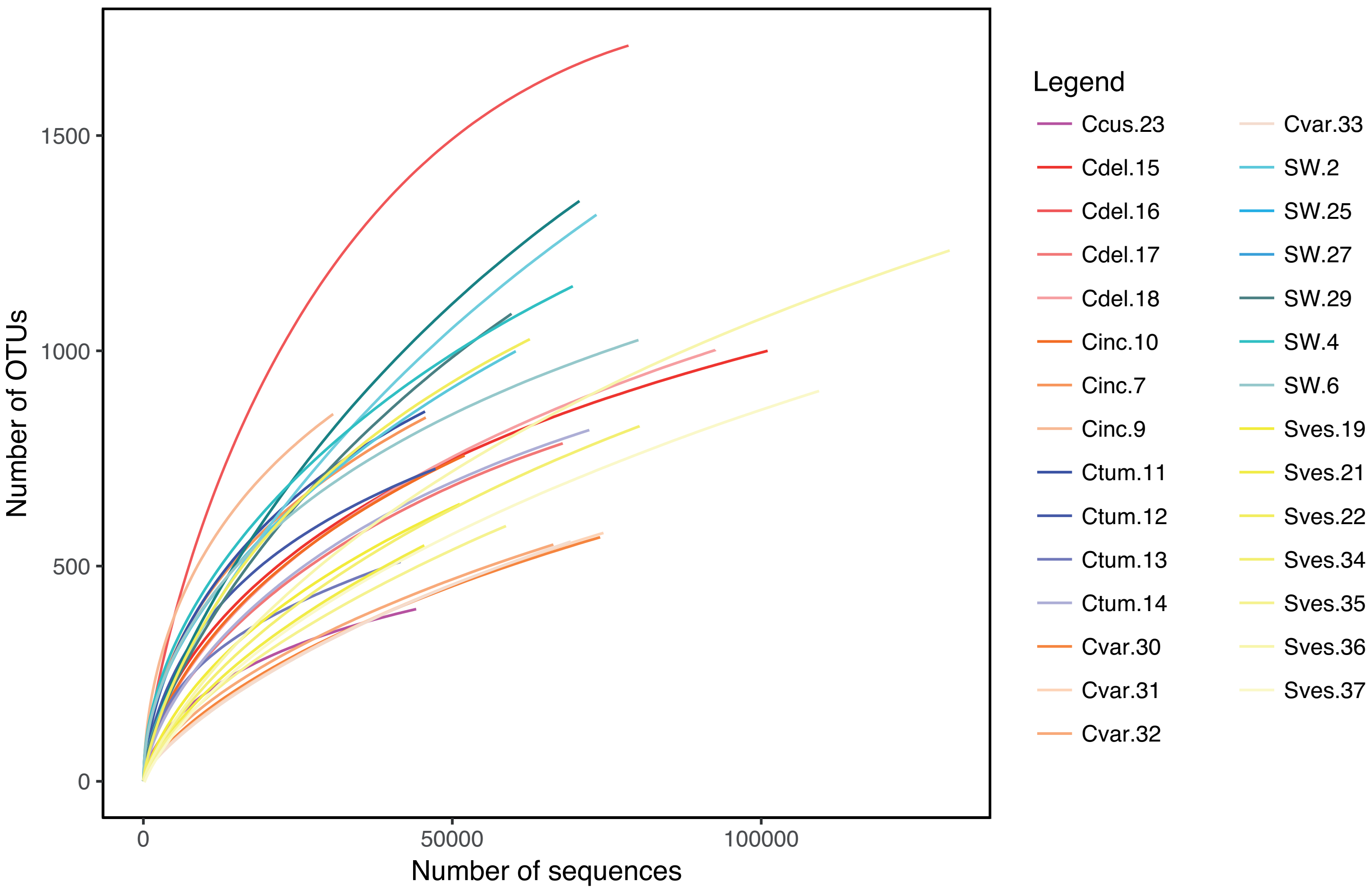

### Supplemental Figure 2

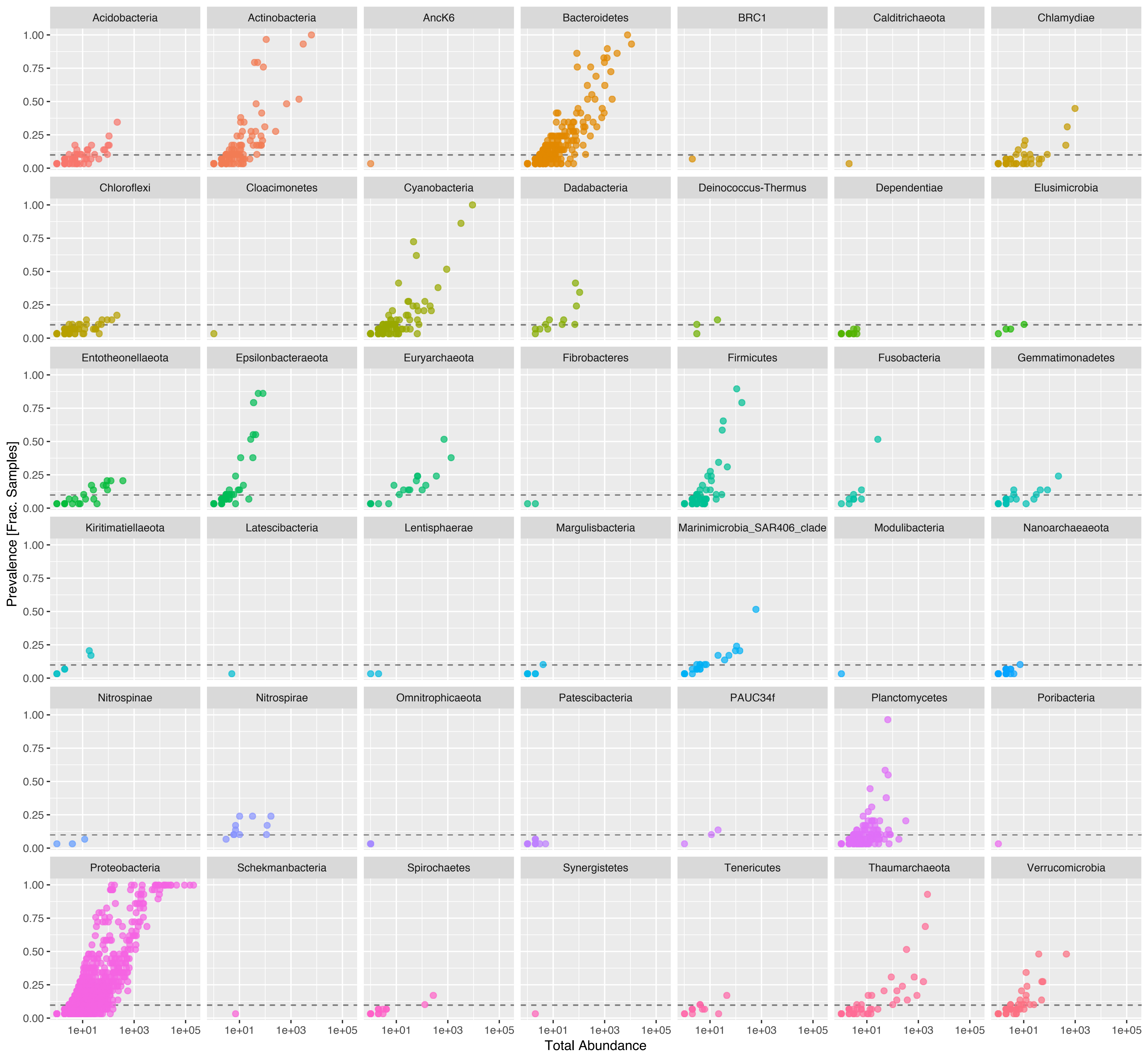

### Supplemental Figure 3

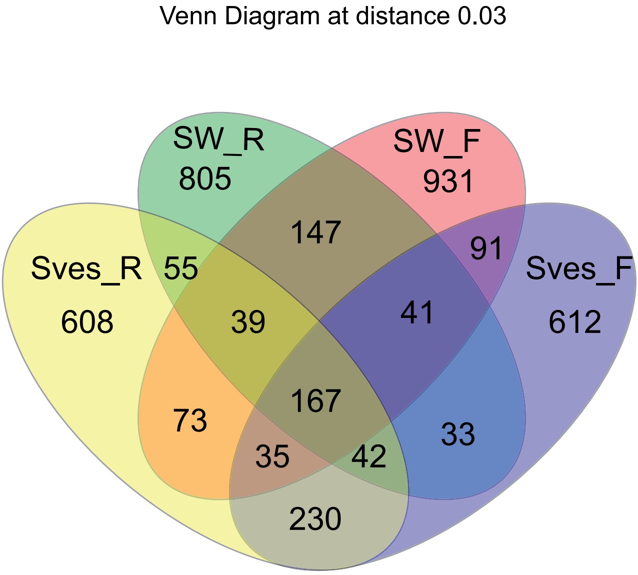

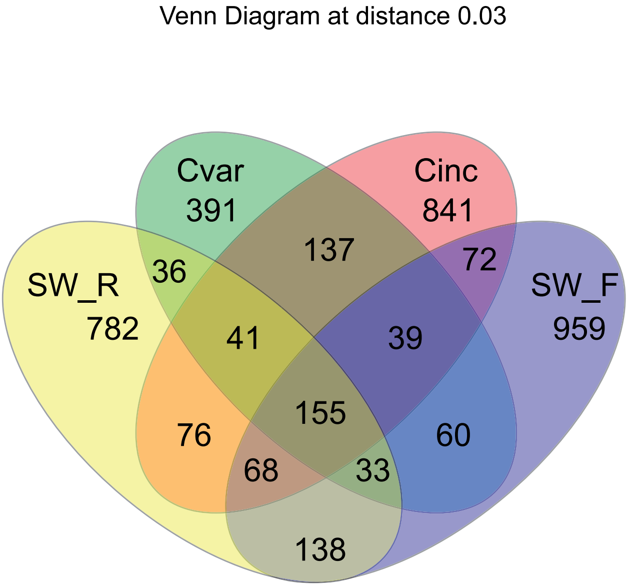

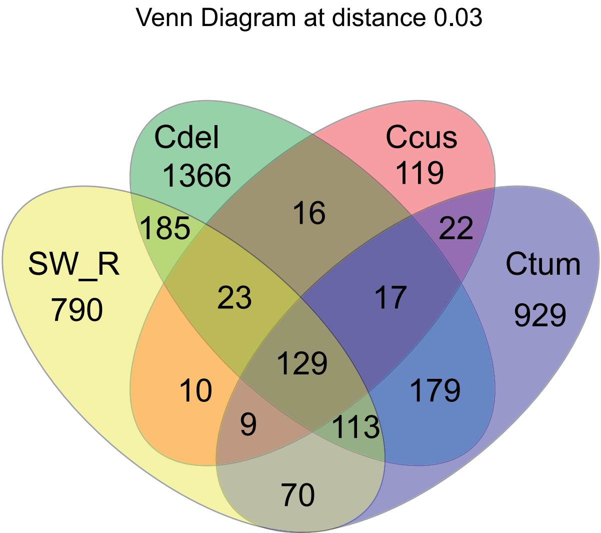

### Supplemental Figure 4

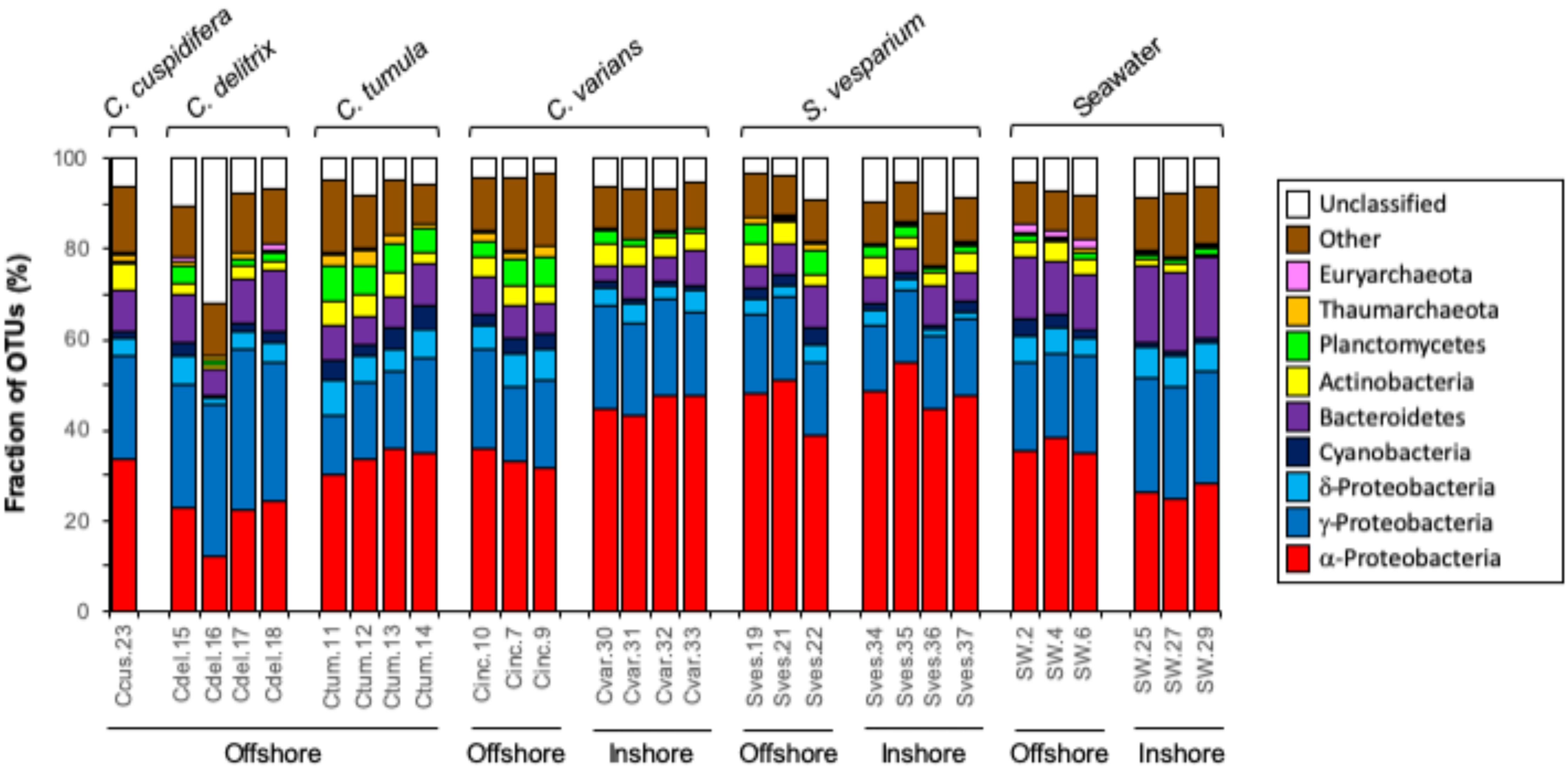

### Supplemental Figure 5

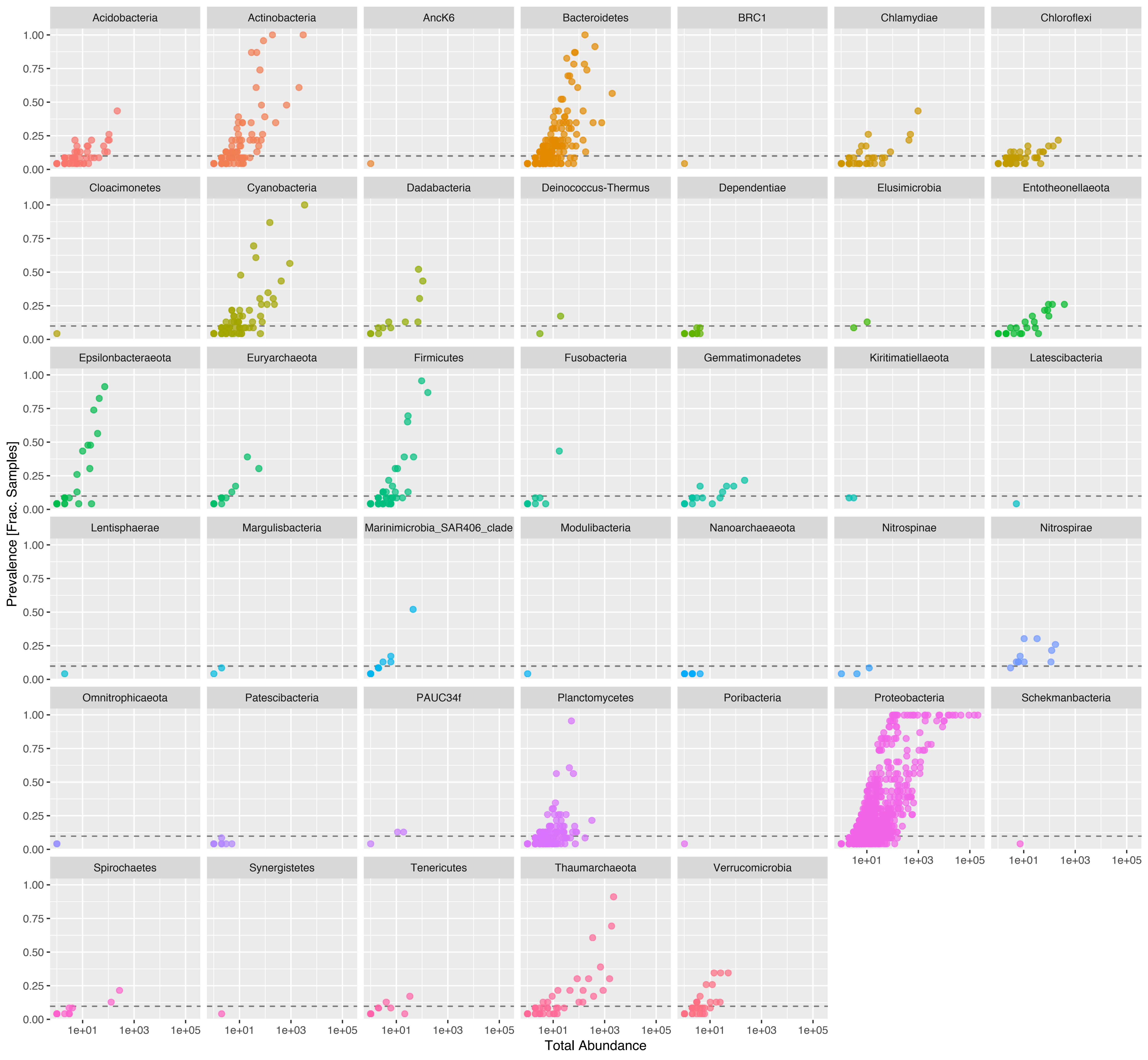

### Supplemental Figure 6

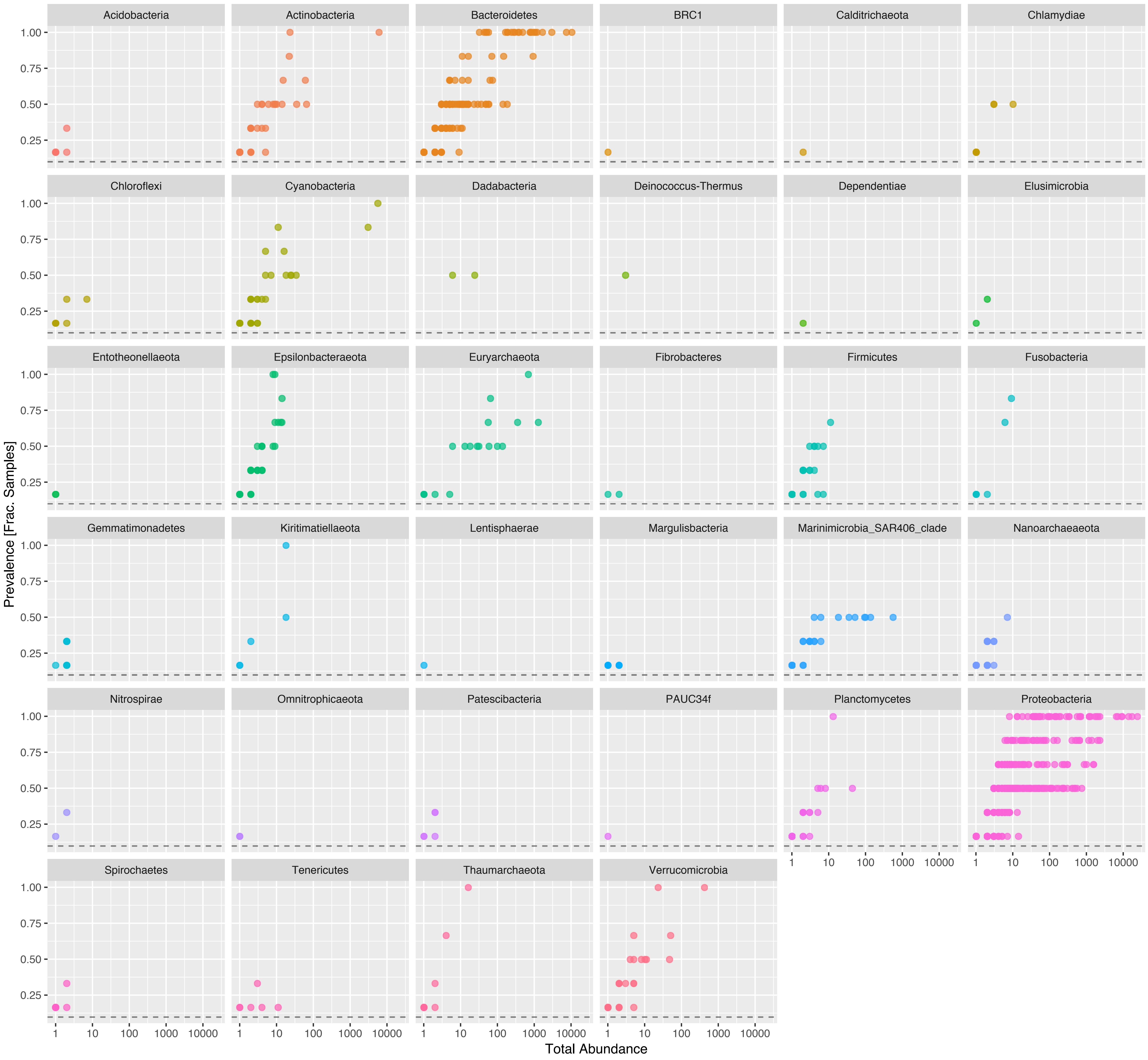
